# Flexibility-aware graph-based algorithm improves antigen epitopes identification

**DOI:** 10.1101/2021.05.17.444445

**Authors:** Chuang Gao, Yiqi Wang, Jie Luo, Ziyi Zhou, Zhiqiang Dong, Liang Zhao

## Abstract

Epitopes of an antigen are the surface residues in the spatial proximity that can be recognized by antibodies. Identifying such residues has shown promising potentiality in vaccine design, drug development and chemotherapy, thus attracting extensive endeavors. Although great efforts have been made, the epitope prediction performance is still unsatisfactory. One possible issue accounting to this poor performance could be the ignorance of structural flexibility of antigens. Flexibility is a natural characteristic of antigens, which has been widely reported. However, this property has never been used by existing models. To this end, we propose a novel flexibility-aware graph-based computational model to identify epitopes. Unlike existing graph-based approaches that take the static structures of antigens as input, we consider all possible variations of the side chains in graph construction. These flexibility-aware graphs, of which the edges are highly enriched, are further partitioned into subgraphs by using a graph clustering algorithm. These clusters are subsequently expanded into larger graphs for detecting overlapping residues between epitopes if exist. Finally, the expanded graphs are classified as epitopes or non-epitopes via a newly designed graph convolutional network. Experimental results show that our flexibility-aware model markedly outperforms existing approaches and promotes the F1-score to 0.656. Comparing to the state-of-the-art, our approach makes an increment of F1-score by 16.3%. Further in-depth analysis demonstrates that the flexibility-aware strategy contributes the most to the improvement. The source codes of the proposed model is freely available at https://github.com/lzhlab/epitope.

**Author summary:** Epitope prediction is helpful to many biomedical applications so that dozens of models have been proposed aiming at improving prediction efficiency and accuracy. However, the performances are still unsatisfactory due to its complicated nature, particularly the noteworthy flexible structures, which makes the precise prediction even more challenging. The existing approaches have overlooked the flexibility during model construction. To this end, we propose a graph model with flexibility heavily involved. Our model is mainly composed of three parts: i) flexibility-aware graph construction; ii) overlapping subgraph clustering; iii) graph convolutional network-based subgraph classification. Experimental results show that our newly proposed model markedly outperforms the existing best ones, making an increment of F1-score by 16.3%.

## Introduction

A B-cell epitope is a specific region at the surface of an antigen that can be neutralized by antibodies, and this neutralization can consequently elicit crucial immune response. Identification of epitopes can be useful to design vaccines, drugs, reagents and so on. Hence, intensive efforts have been made to develop epitope prediction models, including experimental-based and computational-based approaches. Experimental methods, such as X-ray crystallography, nuclear magnetic resonance and phage display [6], are accurate but labor intensive as well as costly, while computational models are more efficient and economical, but suffer from lower accuracy. Due to the difficulty of improving experimental approaches and the fast-evolving computational techniques, massive efforts have been made from the computational perspective.

A general protocol of the computational-based epitope prediction models is: i) collecting antigen-antibody interaction complexes to obtain epitopes; ii) engineering features of epitopes from chemo-physical and statistical perspectives, such as residue’s polarity [7], hydrophobicity [8], protrusion index [9], relative frequency [10], et al.;iii) building computational models based on the features, such as machine learning models [11], graph models [12], statistical models [13], et al.; iv) identifying new epitopes via the well trained models. Although dozens of methods have been proposed, the prediction accuracy still has a big room to improve. For instance, the F1-score of the state-of-the-art epitope prediction model is still less than 0.6 [6]. One possible reason could be the ignorance of structural flexibility of antigens. To our best knowledge, all existing approaches, either experimental or computational, are based on the static structures of antibody-antigen interacting complexes, that is, the structures of antigens and antibodies are fixed when epitopes are determined. However, the structure of an antigen can be flexible [14, 15], particularly the side chain of the surface residues. Therefore, incorporating flexibility into epitope prediction models could be helpful and promising.

Protein flexibility has been widely reported, and there are three types of flexibility, i.e., local flexibility (at atom/residue level) [16], regional flexibility (at intra-domain/multi-residue level) [17], and global flexibility (at multi-domain level) [18]. The local and regional flexibility are mainly caused by the movement of chemical bonds and bond angles [19], while the global flexibility mostly comes from the hinges, helical-to-extended conformations and side chains. Taking as an example shown in Fig 1, the two structures (having the Protein Data Bank (PDB) [23] identifier of 1A14 and 7NN9, respectively) have exactly the same sequence of residues, but notably different structures after aligned. The 1A14 is a bonded structure interacting with the anti-influenza virus neuraminidase, while the 7NN9 is an unbound structure. After alignment, the averaged RMSD (root mean square deviation) of the atoms between the two structures is 0.434 *±* 0.636Å, while the RMSD of the atoms at the epitope region is 0.576 *±* 0.748Å, which is 1.33 times larger than the former one, indicating a markedly movement of conformation upon binding. More interestingly, the side chain RMSD of the epitope residues is 0.853 *±*0.993Å, showing a big fluctuation; cf. Fig 1(A) and (C). These observations indicate that using bonded epitope conformations as training data to figure out epitopes from antigens that have unknown binding partners, to some extent, is inaccurate. This has also been argued by Zhao et al [6]. Unfortunately, all existing models, both experimental and computational, have overlooked this property.

**Fig 1.**
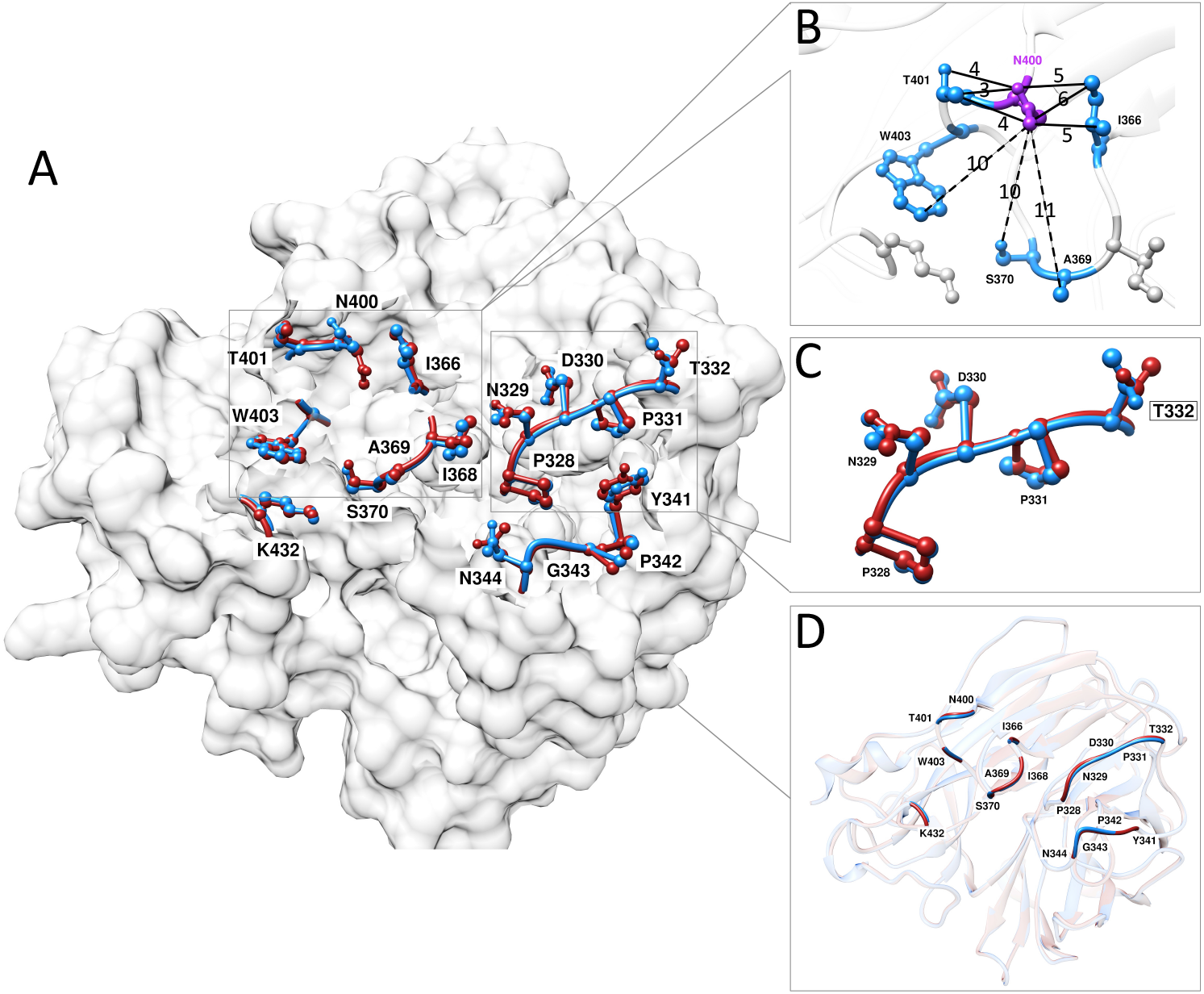
Flexibility illustration. Panel A shows the superimposed structures of a neuraminidase from an influenza virus with (PDB ID 1A14) and without (PDB ID 7NN9) associated antibodies, in which the epitopes are highlighted in balls and sticks. Panel B shows the connections between the residue asparagine (on chain N of 1A14 at position 400) and its neighbors. The solid lines are the connections without considering flexibility, while the dash lines are the connections after flexibility is incorporated. The number on the lines are the distance between the linked atoms measured in angstrom (Å) based on the complex of 1A14. Panel C illustrates the spatial discrepancy between the epitopes on 1A14 and 7NN9, and Panel D shows the superimposed structure between the two in cartoon without side chains. Structures are downloaded from the Protein Data Bank [23] and the figures are generated using the Chimera [24].

To overcome this problem, we propose a graph model that fully considers the local flexibility for epitope prediction; see Fig 2. This model starts with flexibility-aware graphs construction by exploring the distortion of the side chain of all surface residues. These edge-enriched graphs are subsequently partitioned into non-overlapping subgraphs by a message flowing algorithm and expanded into overlapping subgraphs via a community detection algorithm. Finally, a graph convolutional network (GCN) is built to discriminate the expanded subgraphs into epitopes or non-epitopes. Experimental results show that the proposed model elevates the F1-score to 0.656, making an increment of 16.3% compared to the state-of-the-art. In addition, this model is able to identify all single, multiple and overlapping epitopes simultaneously.

**Fig 2.**
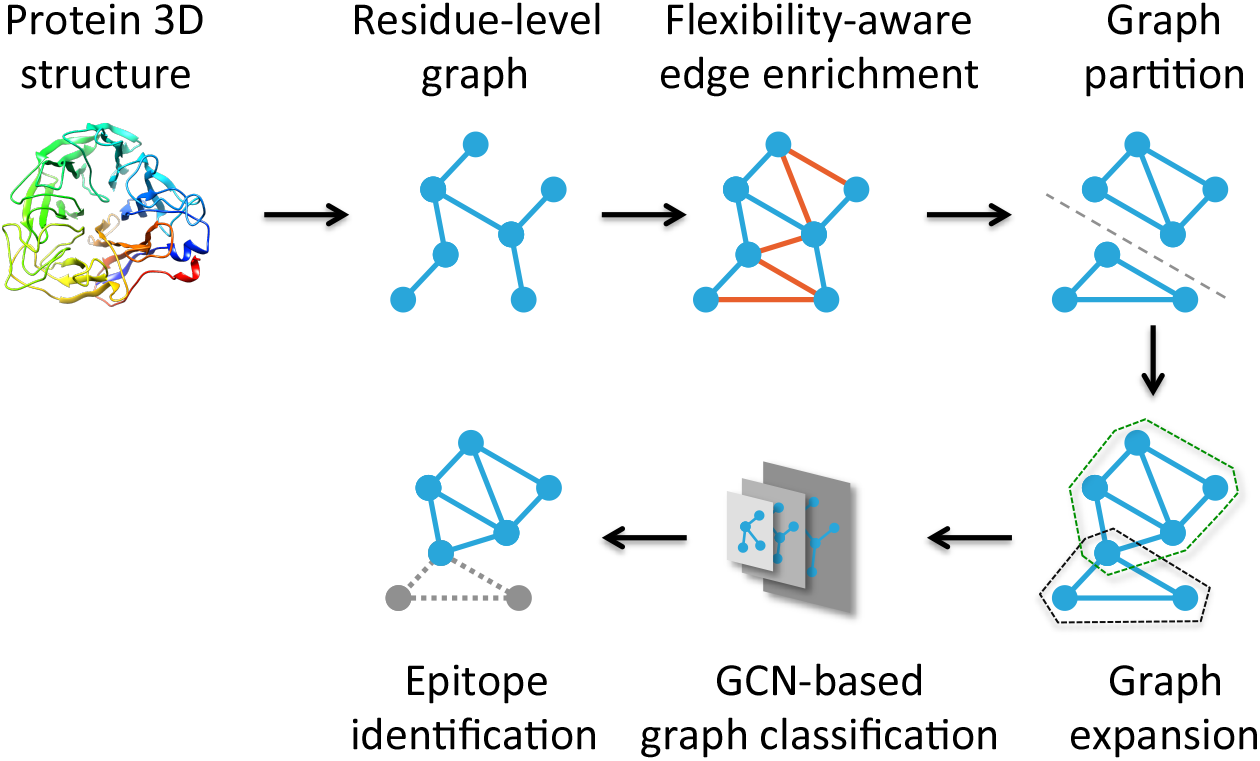
The diagram of the proposed flexibility-aware epitope identification model. It contains graph construction from antigen tertiary structures, flexibility-aware graph enrichment, edge-enriched graph partition, subgraph expansion and graph convolutional network-based classification. The core idea is the inclusion of flexibility in graph construction, which enables big improvement on all the downstream processes.

## Materials and methods

### Data collection

The antigen-antibody interacting complexes are obtained from published data [6], in which all redundant antigens as well as epitopes are removed, and single, multiple, even overlapping epitopes have been well recorded. This dataset contains 258 antigen-antibody complexes, in which 163 antigens have only single epitope (each antigen has only one epitope), 42 have multiple epitopes (each antigen has more than one separated epitopes) and 53 have overlapping epitopes (each antigen has at least one pair of epitopes that are overlapped with each other).

### Flexibility calculation

The conformation of the composing residues of an antigen are not fixed, rather they are quite flexible. This flexibility mainly comes from the rotation of substructures around the covalent bonds [27], particularly the local flexibility (at atom/residue level) [16] and regional flexibility (at intra-domain/multi-residue level) [17], which can be quantified by the torsion angles of residues [28–30]. See for the detailed torsion angles of the twenty standard amino acids.

A residue can have several torsion angles based on the number of atoms located at the side chain. For instance, Alanine has no torsion angle, while Proline has five torsion angles; cf. Table S1. A torsion angle is determined by a tuple of four consecutive atoms within a residue [31], in which the angle is the dihedral angle between the two planes formed by three continuously connected atoms. Take the tuple (C, C_*α*_, C_*β*_, C_*β*2_) in Fig 3 as an example, the torsion angle *χ*_1_ can be considered as the angle between the plane (C, C_*α*_, C_*β*_) and (C_*α*_, C_*β*_, C_*β*2_). All torsion angles are determined analogously by a sliding a widow of four atoms along a residue.

**Fig 3.**
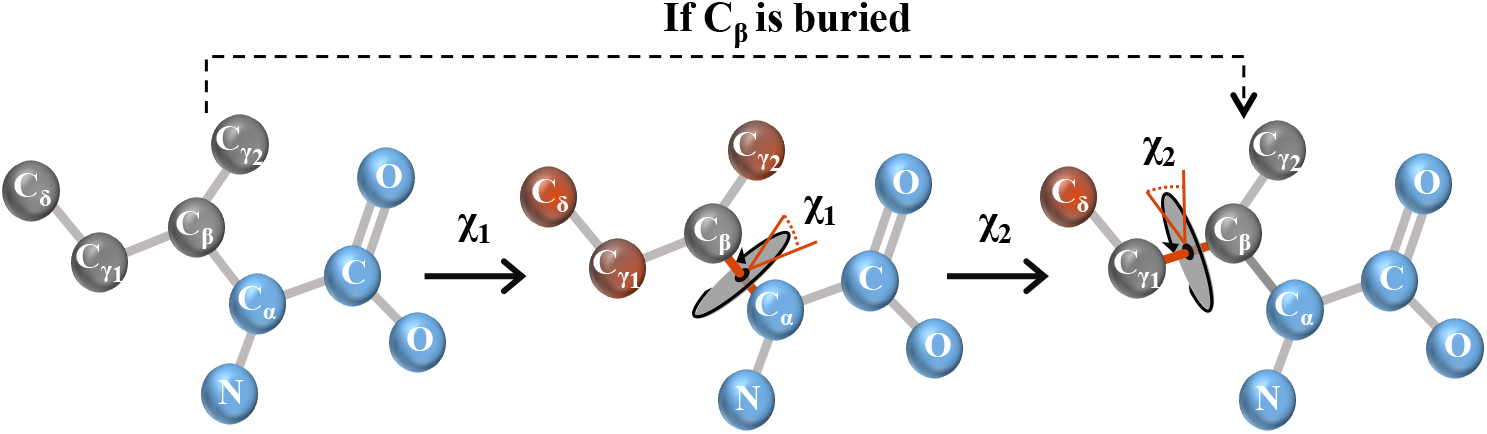
Flexibility calculation illustrated by isoleucine. The possible locations of the atoms at the side chain of the residue are determined by rotating the side chain around axles one-by-one from the C_*α*_ to the farthest atom C_*δ*_. The backbone atoms that are almost static are shown in blue, while the side chain atoms are in gray. The atoms to be affected by rotating the side chain around an axle are shown in red. In case C_*β*_ is buried, we will skip the torsion of *χ*_1_.

To compute the oscillation of each atom due to distortion, we build a tree structure in which the root node is the *C*_*α*_ atom, the leaf node is the farthest atom, and the intermediate nodes are the atoms between the two. Note that, there can have multiple leaf nodes; cf. Fig 3. The content contained in each node is the trajectory of the atom after distorted. Suppose the vector of an arbitrary atom is **v** = ⟨ *x, y, z*⟩ and the unit vector of the rotation axis is **k**, then the rotated vector **v**_*rot*_ of **v** is

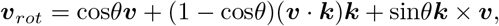

where *θ* is the angle of the rotation. By this means, we are able to determine the possible spatial locations of each atom from its static structure after distortion.

Note that, the flexibility of a buried atom is markedly reduced, even to none, after protein folding. Thus, we only calculate the flexibility of the surface atoms in this study. That is, the root node is replaced by the most superficially buried atom that is connecting with the exposed atom during flexibility determination; cf. Fig 3.

### Model construction

The proposed model is mainly composed of three steps: (i) flexibility-aware graph construction, (ii) overlapping subgraph clustering and (iii) graph classification. The details of each step are as follows.

#### Step 1: flexibility-aware graph construction

A graph *G* = (*V, E*) is generated from the surface atoms of an antigen, where *V* is the set of nodes (surface atoms) and *E* is the set of edges (connections) between atoms in *V*. An atom is deemed as a surface atom if its accessible surface area is no less than 10Å^2^ [12] calculated by NACCESS [32] with the default probe size, while an edge is generated if the Euclidean distance between any two nodes is less than 8Å [12].

Unlike existing approaches of building the graph from a static structure of an antigen (*a snapshot of huge number of possible states*), we consider its flexibility, and generate additional edges for the graph by distortion; see the section ‘Flexibility calculation’. By this means the edges of the graphs can be highly enriched; see Fig 1(B). For instance, after enrichment on edges of the graph generated from PDB 1A14, the number of edges is increased from 635 to 936. To accelerate the determination of edges, the KD-tree data structure is borrowed to calculate the distance between atoms.

The atom-level graphs are upgraded into residue-level graphs by removing redundant and self-contained edges. An edge is deemed as redundant if there exist more than one edge connecting the two nodes, while the self-contained edges are the ones generated from the atoms of the same residue.

#### Step 2: overlapping subgraph clustering

The residue-level graph is clustered into overlapping subgraphs via two steps: partition and expansion. Partition splits the whole graph into non-overlapping subgraphs by using the Markov clustering algorithm [25], while the expansion enlarges separated subgraphs into overlapping subgraphs by using the local community detection algorithm DMF-R [26].

Graph partition is conducted on the weighted graph by using MCL [25] with default parameters, in which the weight *w*_*i,j*_ of an edge *e*_*i,j*_ is calculated as

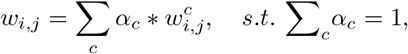

where

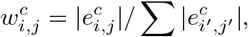

*i, i*^*′*^,*j* and *j*^*′*^ belong to the twenty types of amino acids, *c* is the type of an edge from ‘epitope’, ‘non-epitope’ or ‘boundary’, and |*x*| denotes the size of *x*. Note, boundary edges should be cut off in reality, thus 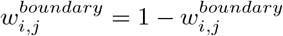.

The ground truth label of an edge is determined from the aforementioned dataset, where an edge 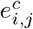 is labed as ‘epitope’ if and only if both the node *i* and *j* are epitopic residues. Similarly, 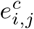 is marked as ‘non-epitope’ if both the node *i* and *j* are non-epitopic residues. An edge 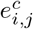 is deemed as a boundary edge if one node is epitopic and the other is non-epitopic.

Subgraph expansion is carried out on the partitioned subgraphs by using the DMF-R algorithm. Unlike existing community detection algorithms that require global information, DMF-R only takes the local community as input and considers its characteristics as well. Hence, it is suitable to our subgraph expansion problem, which is in line with the nature of spatially proximity of epitopes. This step is critical to identify overlapping epitopes as they have been widely reported [33, 34] while graph partition cannot solve this problem intrinsically. To our knowledge, there is only one study aiming at overlapping epitope identification [6].

Note that, the quality of graph partition and expansion are highly related to the way of graph construction, i.e., flexibility-aware or flexibility-agnostic. With flexibility-aware graph construction, hubs may appear; otherwise, all nodes will have evenly distributed degree. See results later.

#### Step 3: graph classification

Expanded overlapping subgraphs are further classified as epitopes or non-epitopes by a newly designed GCN-based algorithm [35, 36], which is composed of two graph convolutional layers and two fully connected layers. The graph convolutional layer accounts for graph embedding, while the fully connected layer is for feature learning.

Let *G* = (*F, A*) be an expanded subgraph having *F ∈ R*^*n×d*^ be the feature matrix and *A ∈ R*^*n×n*^ be the weighted adjacency matrix, where *n* is the number of nodes and *d* is the number of dimensions. The *i*-th graph convolutional layer is calculated as

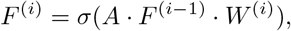

where 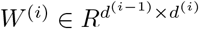 is a trainable weight matrix to be optimized and *σ*(·) is the activation function realized as ReLU [37]. Regarding the output of the *j*-th fully connected layer, it is computed as

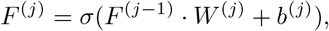

where 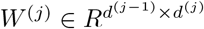 is the parameter matrix to be learned, *b*^(*j*)^ *∈ R*^*n×*1^ is the bias and *σ*(·) is ReLU as well. All the weights *W* ^(·)^ are optimized by the stochastic gradient descent algorithm.

Due to the imbalanced nature of epitopic and non-epitopic subgraphs, we use the focal loss to optimize the training process, which is defined as

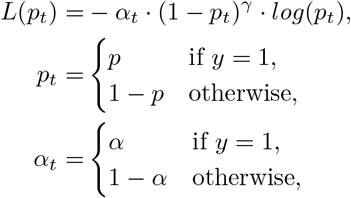

where *y ∈ {*0, 1*}, p ∈* [0, 1], *α∈* [0, 1] and *γ* equals 4.

## Experimental results

### Evaluation metrics

The F1-score, recall and precision are used to quantify the performance of our model as well as existing models, which are defined as:

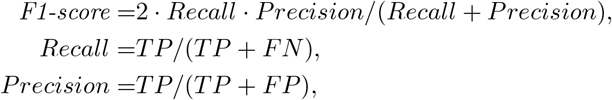

where TP (True Positive) is the number of epitopes fished out correctly, FP (False Positive) is the number of non-epitopes predicted as epitopes incorrectly, and FN (False Negative) is the number of epitopes deemed as non-epitopes. Among these metrics, F1-score is more robust and meaningful as the number of non-epitopes is far larger than that of epitopes.

### Performance qualification

Our proposed model is evaluated by using leave-one-out cross validation on the 258 complexes. The averaged F1-score, recall and precision are 0.656 *±*0.138, 0.587*±* 0.191 and 0.818 *±*0.150, respectively. Comparing to the exiting best mode Glep [6], the proposed model has made remarkable advancement on epitope prediction, achieving an increment of F1-score by 16.3%. On the same dataset, the F1-score achieved by existing approach ElliPro [39], DiscoTope 2.0 [40], EpiPred [41] and Glep [6] is 0.372, 0.159, 0.353, and 0.564, respectively. The detailed performances are shown in Table 1.

**Table 1.**
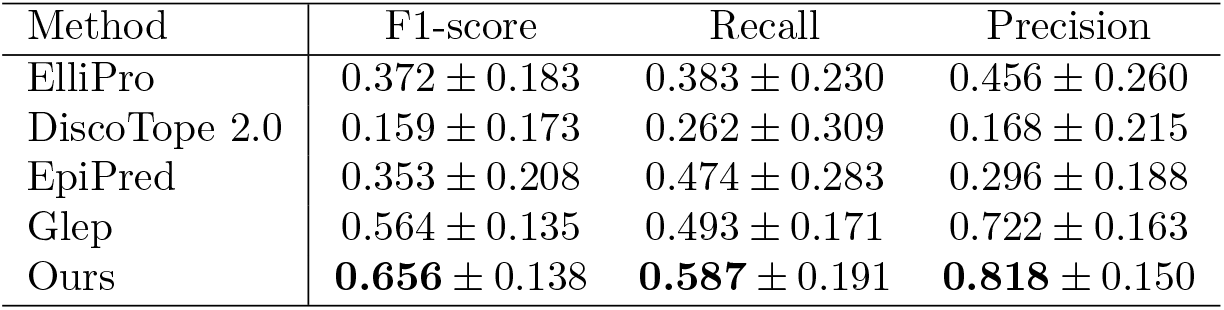
Performance comparison of epitope prediction between our method and the existing state-of-the-art models.

From the table we can see that the performance of our model significantly outperforms existing models. In terms of F1-score, the increment is 76.3%, 312.6%, 85.8% and 16.3% compared with ElliPro, DiscoTope 2.0, EpiPred and Glep, respectively. Besides, the recall and precision obtained by our model are markedly better than that of others as well. The performances of other models are generated from their source codes with default parameters.

### Flexibility enables performance advancement

The key contribution of this study is the incorporation of flexibility into the epitope prediction model. Thus, we carefully examine the impact of flexibility by keeping all other steps unchanged but toggling flexibility only. On average, the F1-score is 0.656*±* 0.138 and 0.599*±* 0.157 for the flexibility-aware and flexibility-agnostic model, respectively. Comparing to the later, the former one lifts the F1-score by 9.5%. Without flexibility the F1-score of the proposed model is only increased by 0.035 compared to the state-of-the-art models [6]. However, this value is increased to 0.092 in case flexibility is considered, which is around 3 times higher. See Fig 4 for the detailed performance comparison.

**Fig 4.**
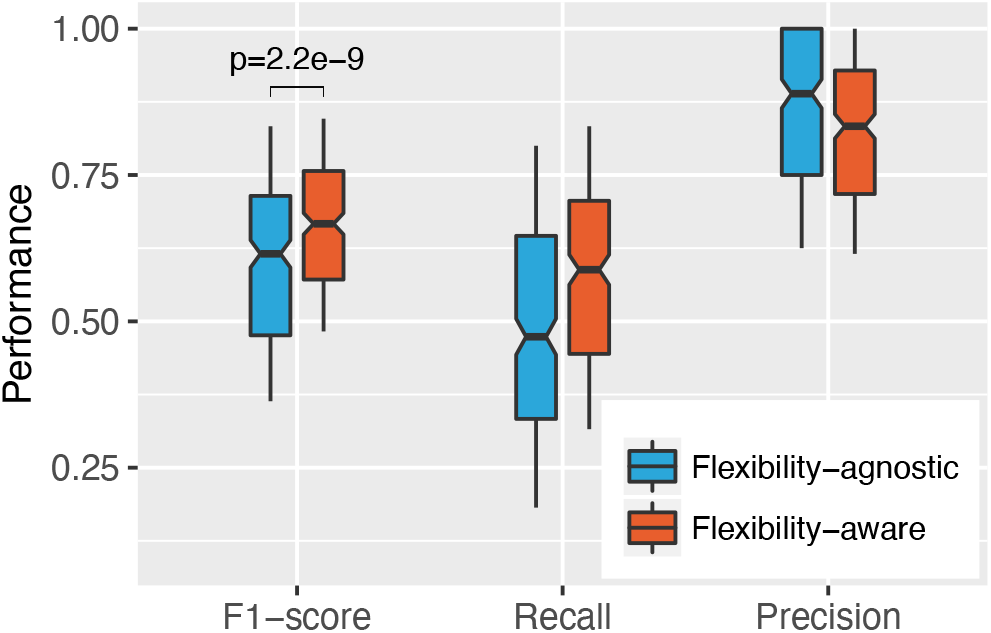
The performance comparison between the flexibility-agnostic and flexibility-aware model.

It is obvious that flexibility has great impact on epitope prediction. We speculate that the impact is mainly from the denser connections of the graphs when flexibility is considered. Based on the data, we found that the averaged degree of each node (residue) was increased from 6.39*±* 1.71 to 9.04*±*2.58, achieving an increment of 41.5%; see Fig 5. Interestingly, the edge enrichment favors nodes with medial degrees, which can be observed from the peak of the bottom left panel of Fig 5. For instance, the averaged difference between the top ten nodes having largest discrepancy of degree is 16.6*±* 0.8, in which the averaged degree of these nodes is 4.9 *±*1.2 and 21.5*±* 0.7 for flexibility-agnostic and flexibility-aware graph, respectively. Notably, these nodes are mainly of Arginine and Lysine, which have been reported as epitope-favored residues [12], indicating that flexibility is particularly helpful to identify epitope-enriched residues. From Fig (the top right panel) we can also see that flexibility enables hubs generation. Hubs are very helpful in graph partition [42], expansion [43] and classification. Without flexibility, the edge-enriched graphs are degradated into ordinary graphs, in which many essential patterns will disappear.

**Fig 5.**
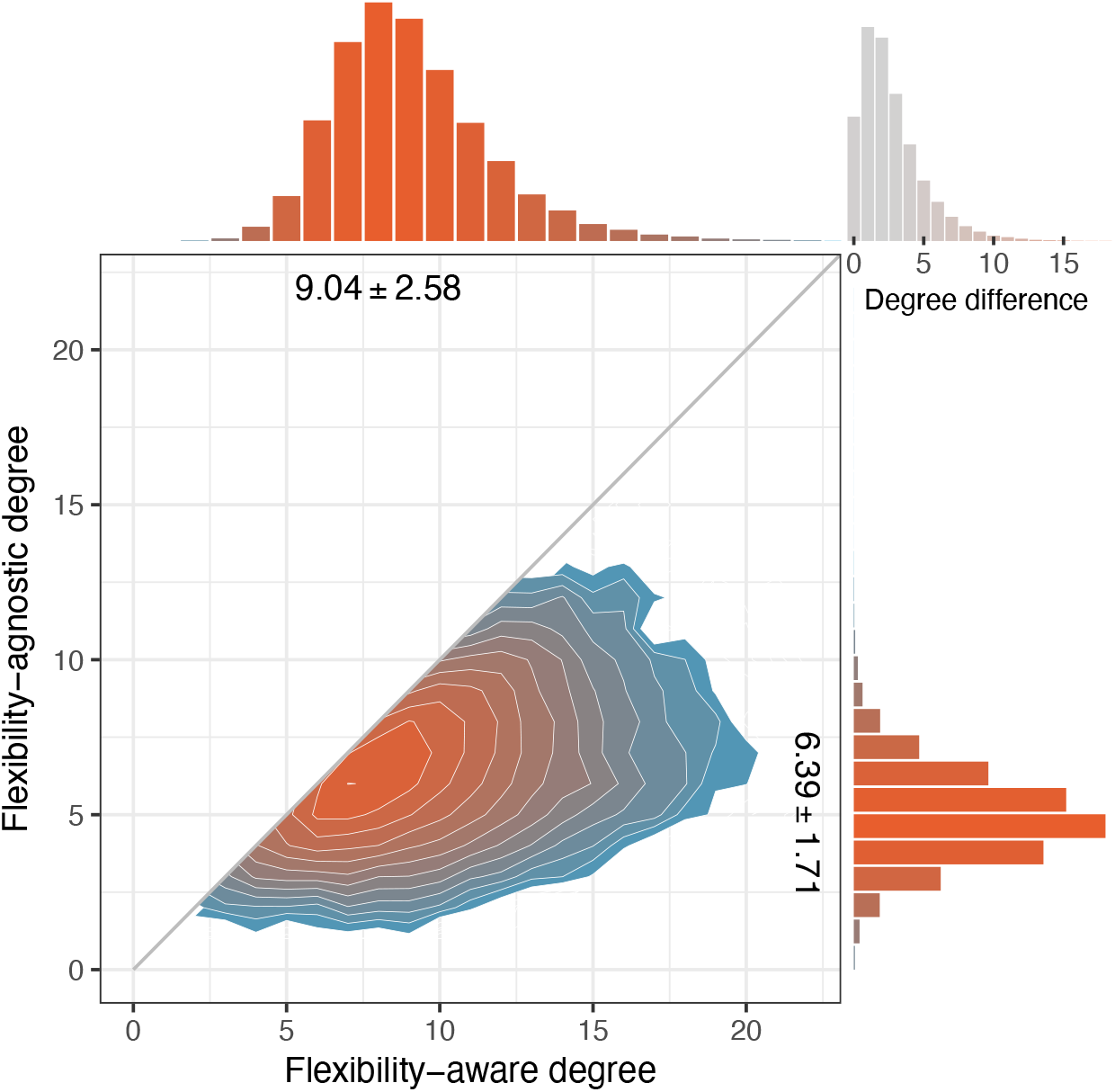
Degree distribution of flexibility-aware and flexibility-agnostic graphs. The color indicates the density of the distribution, of which orange is denser. The degree difference is determined on the same residue with and without flexibility being considered.

### Graph expansion identifies various epitopes

Most existing epitope prediction models are actually epitopic-residue oriented although they are claimed as epitope oriented. This is because the interaction between antigens and antibodies is not necessarily a one-to-one correspondence. In fact, an antigen can interact with multiple antibodies simultaneously or competitively, resulting in several epitopes either separated or overlapped [44, 45]. These epitopes together, however, do not form a big epitope, rather a set of epitopic residues. To solve this problem, we expand subgraphs that are generated from graph partition into larger ones so that shared epitopic residues can be included by more than one epitope, which facilitates the identification of overlapping epitopes.

Among the dataset, there have 53 overlapping epitopes. By using our approach, we can identify 96.2% of them, achieving an averaged F1-score of 0.678 at residue level. Compared to the unexpanded version, the averaged F1-score is increased by 31.6%.

Expansion is also helpful to improve the performance of single and multiple epitopes identification; see Fig 6. Results show that expansion lifts the averaged F1-score by 23.3% and 27.9% for single and multiple epitopes, respectively. Expansion is therefore more helpful for multiple, and overlapping epitopes prediction, particularly the overlapping ones. The F1-score is 0.643, 0.678 and 0.678 for single, multiple and overlapping epitopes identification, respectively; see Fig 6.

**Fig 6.**
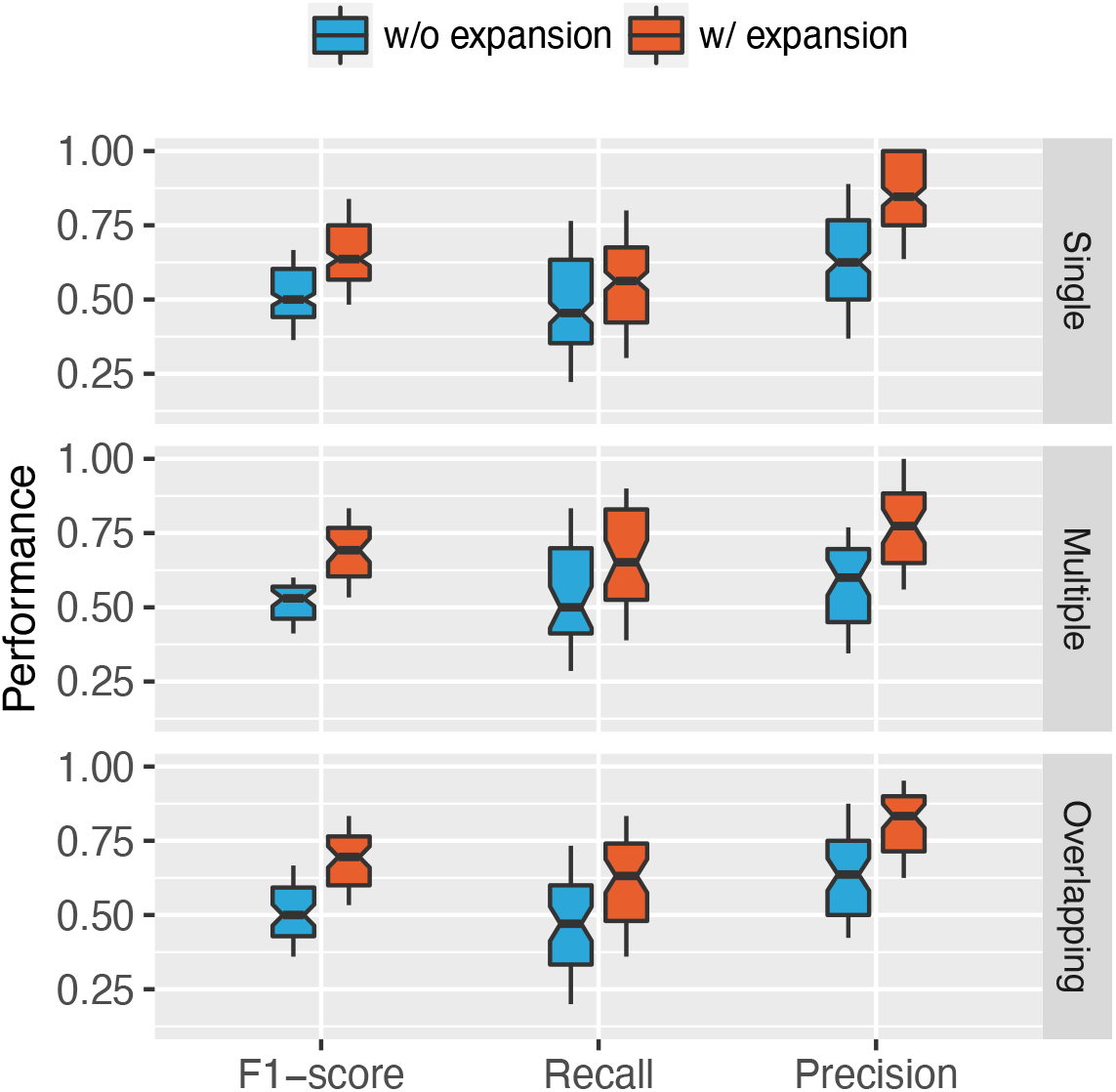
Prediction performance comparison on graphs with and without expansion. The results are broken down into three categories according to the types of epitopes, i.e., single, multiple and overlapping epitopes, to examine the effectiveness of expansion under various circumstances.

### GCN guarantees good distinguishability

The expanded subgraphs are finally classified as epitopes or non-epitopes by using a GCN-based model. The parameters of this model are optimized based on 3,289 subgraphs, in which 935 are deemed as epitopes and the rest as non-epitopes. Since graph partition and subgraph expansion are not perfect, some residues within an epitopic graph are not necessarily epitopic residues. Hence, we consider a subgraph as an epitope if the proportion of epitopic residues is no less than 30% during model training.

The training and validation loss as well as the classification accuracy is show in Fig 7. After two rounds of learning rate decrement, the model is converged to a stable minimum. At this state the best classification accuracy of the test set is 83.1%, and the corresponding F1-score is 0.704.

**Fig 7.**
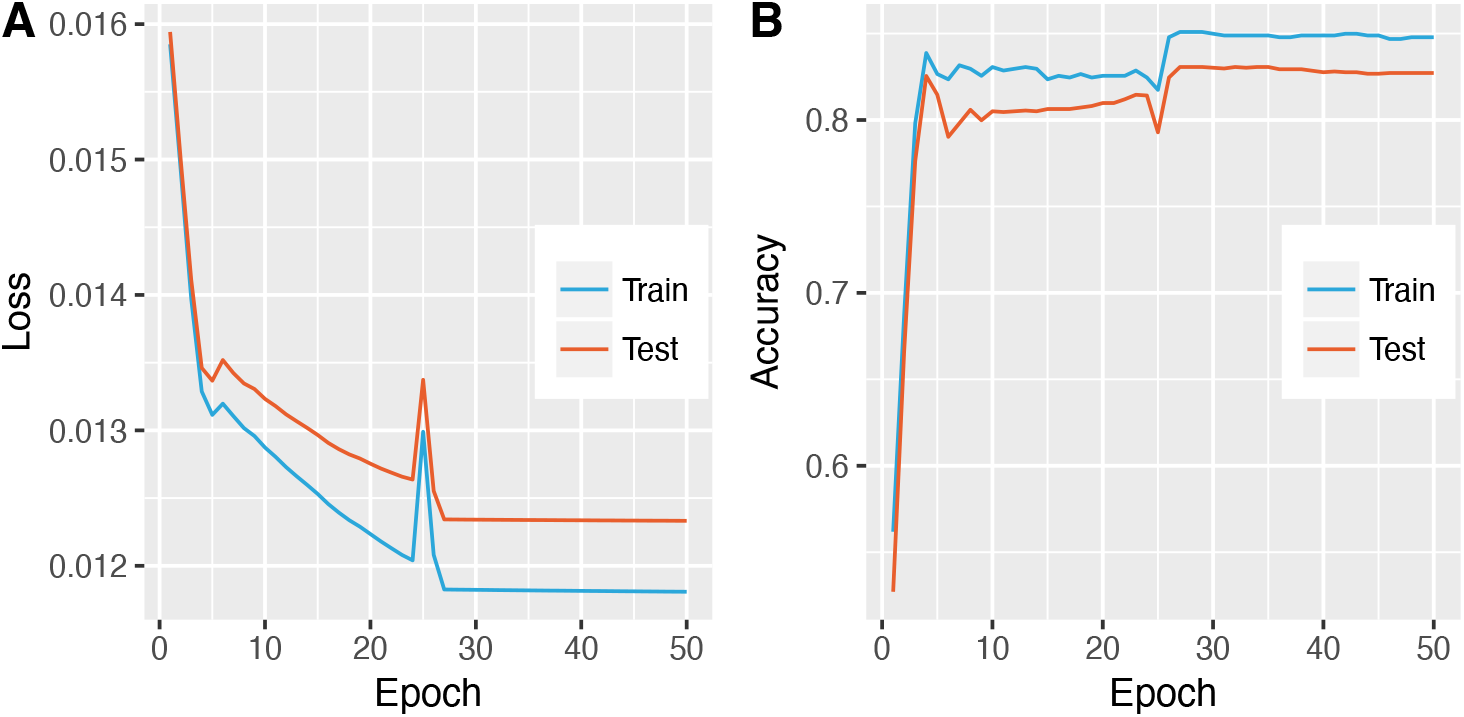
The loss and accuracy of the GCN-based epitope classifier. The model is trained with manually declined learning rate starting from 0.01, and reduced by a degree of 10 in case the loss is unchanged.

### Concluding remarks

Epitopes of an antigen can be neutralized by antibodies, identification of these epitopes can be helpful to various applications, e.g., vaccine design and drug development. Hence, intensive efforts have been made to solve this thorny problem, from both experimental [46, 47] and computational perspectives [48, 49], particularly the later one. Although chemo-physical properties of antigens, such as hydropathy index [50], protrusion index [39] and statistical proximity [51], have been well explored to identify epitopes, all of them are determined from the static structure of antigens, either bonded or unbound. However, the structure of an antigen is not fixed, particularly the side chain of the surface residues [52]. To incorporate such important information, we have proposed a novel graph-based model that is flexibility-aware. Our model starts with residue-level graphs construction from antigens having tertiary structures, and enriches the edges of the graphs via side chain distortion. These edge-enriched graphs are further partitioned into subgraphs by using MCL [25], and are expanded into overlapping subgraphs by DMF-R [26]. Finally, these expanded graphs are classified into epitopes or non-epitopes by a newly built graph convolutional network. Experimental results show that the proposed approach outperforms all existing models, achieving a high F1-score of 0.656. This score makes a 16.3% increment compared to that of the state-of-the-art models. Besides, our model can identify single, multiple and overlapping epitopes simultaneously. Although there still has notably room to improve the performance, this study pioneers a new direction for improving epitopes identification.

## Supporting information

**Table S1.** The torsion angles of the twenty standard amino acids.

| Amino acid | Torsion angle ($^{\circ}$) | | | | | Reference |
| --- | --- | --- | --- | --- | --- | --- |
| | $\chi_1$ | $\chi_2$ | $\chi_3$ | $\chi_4$ | $\chi_5$ | |
| GLY | — | — | — | — | — | — |
| ALA | — | — | — | — | — | — |
| CYS | — | — | — | — | — | [1] |
| VAL | 21 | — | — | — | — | [1] |
| SER | 16 | — | — | — | — | [1] |
| THR | 16 | — | — | — | — | [1] |
| ILE | 45 | 47 | — | — | — | [2] |
| LEU | 36 | 40 | — | — | — | [2] |
| ASP | 29 | 71 | — | — | — | [2] |
| ASN | 21 | 59 | — | — | — | [2] |
| HIS | 22 | 56 | — | — | — | [2] |
| TRP | 30 | 38 | — | — | — | [2] |
| PHE | 29 | 71 | — | — | — | [1, 2] |
| TYR | 17 | 46 | — | — | — | [1, 2] |
| GLU | 30 | 39 | 71 | — | — | [1, 2, 3] |
| GLN | 30 | 39 | 59 | — | — | [1, 2, 3] |
| MET | 36 | 39 | 38 | — | — | [2] |
| LYS | 29 | 36 | 32 | 40 | — | [2] |
| ARG | 31 | 30 | 37 | 40 | 7 | [2] |
| PRO | 26.8 | 36.5 | 32.2 | 17.3 | 9.8 | [4] |

Note, the side chain torsion angle must be formed from at least two heavy atoms, thus the GLY and ALA have no torsion angle.

## Acknowledgments

This study was collectively supported by the Natural Science Foundation of China [32060150; 31871481], the Natural Science Foundation of Guangxi [2018GXNSFAA281275], the Free Exploration Fund of Hubei University of Medicine [FDFR201805], the Scientific Research Found of Guangxi University [XGZ150316] and Taihe Hospital [2016JZ11].

## Notes

### Competing Interest Statement

The authors have declared no competing interest.

